# Evaluation of stability and inactivation methods of SARS-CoV-2 in context of laboratory settings

**DOI:** 10.1101/2020.09.11.292581

**Authors:** Sandra Westhaus, Marek Widera, Holger F. Rabenau, Sebastian Hoehl, Denisa Bojkova, Jindrich Cinatl, Sandra Ciesek

## Abstract

The novel coronavirus SARS-CoV-2 is the causative agent of the acute respiratory disease COVID-19, which has become a global concern due to its rapid spread. Laboratory work with SARS-CoV-2 in a laboratory setting was rated to biosafety level 3 (BSL-3) biocontainment level. However, certain research applications in particular in molecular biology require incomplete denaturation of the proteins, which might cause safety issues handling contaminated samples. In particular, it is critical to provide proof of inactivation before samples can be removed from the BSL-3.

In this study, the stability of the virus in cell culture media at 4°C and on touch panel surfaces used in laboratory environment was analyzed. In addition, we evaluated common lysis buffers that are used in molecular biological laboratories for their ability to inactivate SARS-CoV-2. We have found that guanidine thiocyanate and most of the tested detergent containing lysis buffers were effective in inactivation of SARS-CoV-2, however, the M-PER lysis buffer containing a proprietary detergent failed to inactivate SARS-CoV-2. Furthermore, we compared chemical and non-chemical inactivation methods including ethanol, acetone-methanol mixture, PFA, UV-C light, and heat inactivation.

In conclusion, careful evaluation of the used inactivation methods are required and additional inactivation steps are necessary before removal of lysed viral samples from BSL-3.

## 1. Introduction

The coronavirus disease 2019 (COVID-19) is caused by an infection with the severe acute respiratory syndrome coronavirus 2 (SARS-CoV-2). The origin of SARS-CoV-2 outbreak was initially described in Wuhan, China (Zhu et al., 2020) and rapidly established a worldwide pandemic. Globally, 57.8 million cases 1.3 million deaths in total since the start of the pandemic have been reported by the WHO as published on November, 24^th^ 2020 (WHO, 2020). Numerous experimental studies are currently being carried out, which require appropriate inactivation methods to restrict laboratory born spread among scientists and lab personnel.

SARS-CoV-2 is a spherical beta coronavirus with a size of 120 nm in diameter, which has a lipid envelope. Persistence and inactivation of coronaviruses including the highly pathogenic SARS-CoV and MERS-CoV, which emerged in the last decades, was evaluated in several publications (reviewed in (Kampf et al., 2020a; Kampf et al., 2020b)). Accordingly, a whole series of chemical and physical inactivation methods such as UV radiation, heat inactivation and detergents are assumed to be effective inactivation methods against SARS-CoV-2 (Kratzel et al., 2020b). However since human coronaviruses were shown to remain infectious on inanimate surfaces at room temperature for up to 9 days (reviewed in (Kampf et al., 2020a; Kampf et al., 2020b)), contamination of frequently touched surfaces in laboratory settings are therefore a potential source of viral transmission.

As shown for other enveloped viruses (Fischer et al., 2015; Goo et al., 2016; Muller et al., 2016), it might be assumed SARS-CoV-2 can remain infectious for much longer period of time when stored in liquid milieu at appropriate temperatures. This is the case in laboratory environments in which the condition of the sample has to be partially preserved for the subsequent molecular biological assays.

Quantitative reverse transcriptase polymerase chain reaction (RT-qPCR) is the major detection method for SARS-CoV-2, in which guanidine thiocyanate containing chaotropic buffers are efficient in disrupting viral structures. However, there are applications in particular in molecular biology that require incomplete denaturation of the proteins (e.g. luciferase measurements). Thus, in this study we evaluated common lysis buffers that are used in molecular biological laboratories for their ability to inactivate SARS-CoV-2. Furthermore, we investigated the stability of SARS-CoV-2 in cooled cell culture media and on touch panels on electronic devices. In addition we analyzed the impact of temperatures, physical (UV) and chemical influences on viral infectivity.

## 2. Materials and methods

### 2.1 Cell culture and virus propagation

Caco2 and Vero cells were cultured in Minimum Essential Medium (MEM) supplemented with 10% fetal calf serum (FCS), 100 IU/ml of penicillin and 100 μg/ml of streptomycin. Viral titers of SARS-CoV isolate HK or FFM1 (Drosten et al., 2003) and SARS-CoV-2 isolate (Frankfurt 1, FFM1) (Hoehl et al., 2020; Toptan et al., 2020) were determined by TCID_50_. For virus propagation, Caco-2 or Vero cells were seeded in a 96-well plate and inoculated with SARS-CoV or SARS-CoV-2 using a MOI of 0.1 (Toptan et al., 2020). After 60 min at 37°C and 5% CO_2_ cells were rinsed with PBS, supplemented with fresh medium, and incubated until harvest. Influenza isolates PR8 (H1N1) [ATCC-VR-1469; p1; Titer: 1.1× 10^7^] and Victoria (H3N2) [ATCC-VR-822; p5; Titer: 1.4× 10^7^] were prepared by infecting MDCK cells. Cell free virus aliquots were stored at −80°C. All infectious work was performed under biosafety level 3 (BSL-3) conditions in a BSL-3 facility according to the Committee on Biological Agents (ABAS) and Central Committee for Biological Safety (ZKBS).

### 2.2 Determination of SARS-CoV-2 infectivity by TCID_50_

Infectious titer of SARS-CoV-2 supernatant were determined by an end-point limiting dilution assay as 50% tissue culture infectious dose (TCID_50_/ml) in confluent cells in 96-well microtiter plates. Briefly, Caco-2 were seeded onto a 96-well plate and infected with untreated or treated SARS-CoV-2 containing supernatant in an initial 1:10 dilution (quadruplicates). Titration was performed on 96-well plate with ten serial 1:10 dilutions. Viral titer was determined 48 h post infection as 50% tissue culture infectious dose (TCID_50_) as described by Spearman and Kaerber (Kärber, 1931; Spearman, 1908). To analyze inhibition capacity of different inactivation methods viral titers were normalized to the untreated control and reduction factor was calculated as described elsewhere (Rabenau et al., 2005b). All infection experiments were performed with initial viral titer of 1× 10^6^ TCID_50_/mL if not indicated differently.

### 2.3 Chemical Inactivation of cell culture supernatants

To test different chemical compounds to inactivate SARS-CoV-2, virus containing supernatant was mixed 1:1 or 1:10 with lysis buffers or fixation solutions, respectively. As adapted from previously published procedures (Rabenau et al., 2005a), the mixture was incubated for 10 minutes at room temperature before further processing. Testing different lysis buffers, compound-virus mixture was further diluted 1:1000 to avoid cytotoxic effect before added to the seeded Caco-2 cells. Viral load and inactivation capacity were determined by RT-qPCR and CV staining, respectively. Analyzing different fixation solutions, compound-virus mixture was further diluted 1:100 before added to the seeded Vero cells. Viral titers and inactivation capacity were determined by RT-qPCR and compared to viral control. Before use Western blot (WB) lysis buffer (20 mM TRIS/HCl pH 7.5, 150 mM NaCl, 10 mM NaPPi, 20 mM NaF, 1% Triton-X, 1.9 M glycine, and 250 mM TRIS base) was supplemented with cOmplete™ protease inhibitor cocktail tablets (Roche).

### 2.4 Physical Inactivation of cell culture supernatants

Analyzing physical parameters to inactivate SARS-CoV-2, virus containing supernatant was treated either at different temperatures or with UV-C light or was applied to different surfaces and stored for a distinct period of time. For surface inactivation SARS-CoV-2 was applied to cell culture dishes (polystyrole; plastic), used smartphone display, and used protection film and dried at ambient temperature. Of note, the used surfaces were not cleaned prior to the experiments to maintain an environment close to real-world conditions. Dried virus spots were incubated for 6 h or 5 days at ambient temperature before wiped off with a PBS soaked swab (2 cm^2^) and eluted in culture medium. Testing different temperatures for inactivation of SARS-CoV-2 virus containing supernatant was incubated on a thermo shaker for defined time at 56°C, 60°C or 90°C. In addition, determination of viral load was performed by infecting Vero cells and measuring gene copies/reaction by RT-qPCR. Finally, SARS-CoV-2, SARS-CoV- and Influenza A containing supernatant was applied to cell culture dishes and dried. Dried virus spots were treated with UV light of different light sources. “UVA-Cube” and UV-C LED (Hönle AG, Germany) both contain a UV-C light source (LED Spot 100 IC / HP IC), which was set to E = 8.8 mW/cm² (365 – 460 nm) with a fixed distance of 2 cm. Benchtop UV-light was emitted using a laminar flow (Holton LaminAir, Heto-Holten, Denmark). After irradiating for a defined time spots were eluted in culture medium. Viral load and inactivation capacity were determined by TCID_50_ and CV staining, respectively.

### 2.5 Rotitest Vital viability assay

To analyze the viability of cells treated different compounds to inactivate SARS-CoV-2, the intracellular dehydrogenase activity was tracked using the Rotitest Vital Kit (Carl Roth GmbH + Co. KG, Karlsruhe, Germany). Briefly, cells were seeded in 96-well plates and incubated with 1:1000 dilution of inactivating compound in culture medium in triplicates per sample. After 48 h of incubation at 37°C and 5% CO_2_, Rotitest Vital solution was added and analyzed using a multimode reader at 450 nm.

### 2.6 RT-qPCR analysis and detection of SARS-CoV-2 genomic RNA

For detection of intracellular and extracellular SARS-CoV-2 genomic RNA, primers M-475-F (5’-TGTGACATCAAGGACCTGCC-3’) and M-574-R (5’-CTGAGTCACCTGCTACACGC-3’) were used together with the Fam-BHQ1 dual-labeled probe M-507-P (5’-TGTTGCTACATCACGAACGC-3’) as described previously (Toptan et al., 2020). For normalization to intracellular mRNA level human GAPDH was measured using primers GAPDH-fwd (5’-TGCACCACCAACTGCTTA) and GAPDH-rev (5’-GGATGCAGGGATGATGTTC-3’).

### 2.7 Determination of SARS-CoV-2 inactivation by Crystal Violet Staining

Inactivation of SARS-CoV-2 by chemical and physical methods was (when indicated) determined by Crystal Violet Staining (CV). Therefore, Caco-2 cells were seeded on a 96-well plate infected with untreated or treated SARS-CoV-2 supernatant and cultured for 48 h at 37°C, 5% CO_2_. Cells were fixed after incubation with 3% paraformaldehyde (PFA) for 20 minutes at room temperature before stained with 0.1% CV for another 20 minutes. Staining solution was removed and plates were dried overnight. CV was dissolved with 100% ice-cold methanol and analyzed using a multimode reader at 560 nm. To determine inactivation capacity samples were normalized to untreated control.

### 2.8 Statistical analysis

If not indicated differently all infection experiments were repeated in three independent experiments. Data analysis was performed in Microsoft Excel and GraphPad Prism 6 (GraphPad Software, USA). Statistical significance compared to untreated control was determined using unpaired student’s t-test on non-log-transformed data. Asterisks indicated p-values as * (p < 0.05), ** (p ≤ 0.01) and *** (p ≤ 0.005).

## 3. Results

### 3.1 SARS-CoV-2 is stable for several weeks

To investigate how long SARS-CoV-2 viruses are stable under 4°C storage conditions, we thawed different stock aliquots of a SARS-CoV-2 strain FFM1 (Hoehl et al., 2020; Toptan et al., 2020) (which were cryo-conserved at −80°C) at weekly intervals and stored them at 4°C in cell culture medium for up to 23 weeks. The long-stored virus stocks were then examined for their infectivity using TCID_50_. Remarkably, even after 23 weeks under these storage conditions we found considerable quantities of infectious virus (**Figure 1a**).

**Figure 1.**
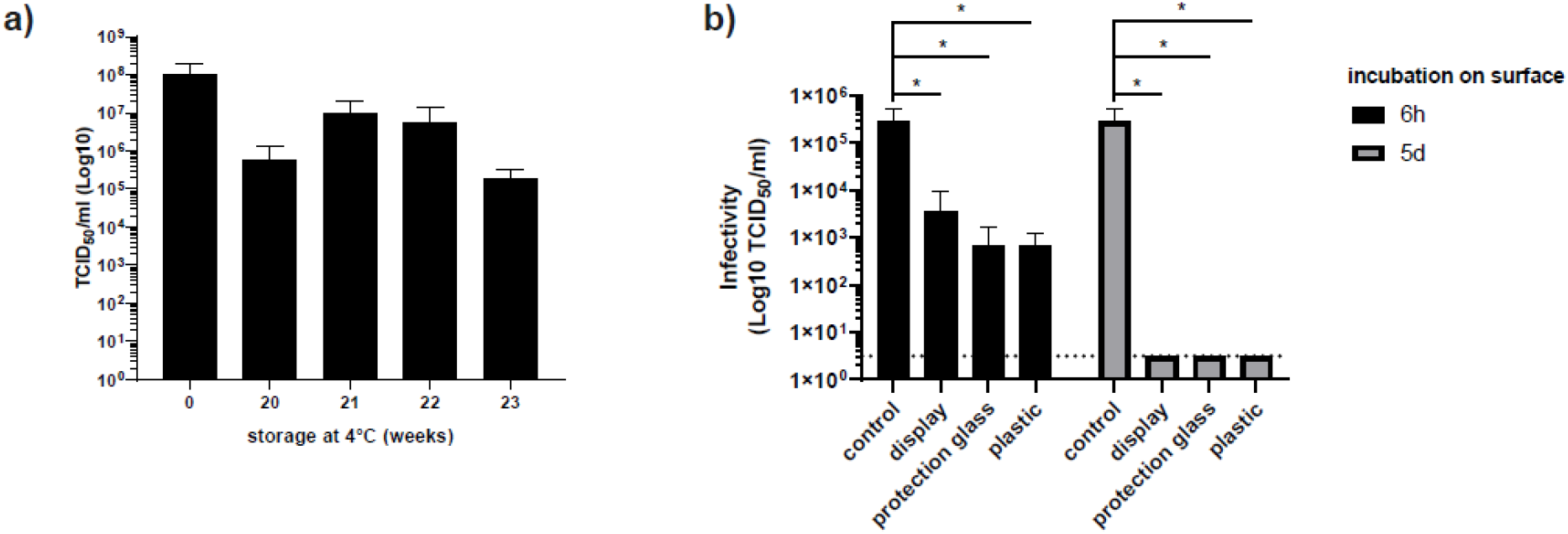
Stability of SARS-CoV-2 under different storage conditions. a) SARS-CoV-2 virus stocks were stored in cell culture medium at 4°C for the indicated period of time. b) Stability of SARS-CoV-2 on surfaces (with and without protection gla*s*s).TCID_50_ using CaCo2-cells was performed to determine viral infectivity (Microscopical CPE readout). Representative experiment performed in quadruplicates. Error bars indicate SD from four technical replicates.

Since coronaviruses were shown to remain infectious on commonly used touch surface materials (Warnes, Little, and Keevil, 2015), we next examined how long SARS-CoV-2 might be stable on touch panels of electronic devices that are commonly used in well-equipped laboratories. For this purpose, we coated plastic as well as touchscreen displays of mobile phones with and without protective glass with SARS-CoV-2 (2.8×10^5^ TCID_50_/mL). After a short and long incubation time at room temperature corresponding to one working day (6 h) and one working week (5 days), the surfaces were wiped off with a PBS soaked swab and the viruses were eluted in culture medium and tested quantitatively for infectivity. After 6 hours we saw a decrease in TCID_50_ of 1.91 log on the display and of approx. 2.6 log for displays with a protective glass and for plastic surfaces (**Figure 1b, and Table 1**). After 5 days we were unable to recover infectious virus particles on any of the surfaces indicating a minimal reduction factor of 4.95 log.

**Table 1.** Stability of SARS-CoV-2 on surfaces in laboratory environment. Common laboratory surfaces plastic and touchscreen displays with and without protection glass were contaminated with infectious SARS-CoV-2. Recovered virus was tested for infectivity performing TCID_50_ assays. Detection limit was 0.5 TCID_50_/mL (Log10).

| Incubation time | Surface | Virus titer<br>(TCID<sub>50</sub>/ml<br>[log10] ± SD) | Minimal<br>reduction factor<br>(log10) | p-value |
| --- | --- | --- | --- | --- |
| Viral control |  | 5.45 ± 5.39 | 0.00 | - |
| 6 hours | display | 3.54 ± 3.75 | 1.91 | 0.0377 |
|  | protection glass | 2.84 ± 2.98 | 2.60 | 0.0357 |
|  | plastic | 2.84 ± 2.74 | 2.61 | 0.0357 |
| 5 days | display | 0.5 ± 0.0 | 4.95 | 0.0352 |
|  | protection glass | 0.5 ± 0.0 | 4.95 | 0.0352 |
|  | plastic | 0.5 ± 0.0 | 4.95 | 0.0352 |

These data indicate that it is particularly important to inactivate contaminated material and instruments before continue working under less stringent safety requirements.

### 3.2 Effectiveness of lysis buffers at inactivating SARS-CoV-2

Molecular biological detection methods require specific lysis methods to ensure subsequent partly enzymatic reactions. Buffers containing guanidine thiocyanate have proven useful for the isolation of viral nucleic acids and PCR-based analysis. We have therefore tested a set of six common lysis buffers for nucleic acid extraction whereof three are also used in routine diagnostic. Buffers AL, ATL and AVL (Qiagen, Hilden, Germany) and the cobas omni buffer (Roche, Germany) are commonly used to isolate nucleic acids from cell culture supernatants and patient material such as serum, blood, plasma, but also throat swab samples. The buffer RLT (Qiagen), however, is mainly used for research purpose for the isolation of total cellular RNA from infected cells. SARS-CoV-2 containing cell culture supernatants were mixed 1:1 with lysis buffers and diluted after 10 minutes of incubation. To evaluate the remaining infectivity viral outgrowth assays were performed in Caco2 cells and after 48 h cell culture supernatants were harvested for analysis. To control that the readout was not affected by cell toxicity issues, in parallel we performed a cell viability assay (**Figure 2a**). In order to monitor virus production, we additionally carried out an RT-qPCR targeting the SARS-CoV-2 M-gene. As shown in **Figure 2** all tested buffers containing guanidine isothiocyanate including the cobas omni buffer (Roche) were able to completely inactivate samples containing SARS-CoV-2 (**Table 2**).

**Table 2.**
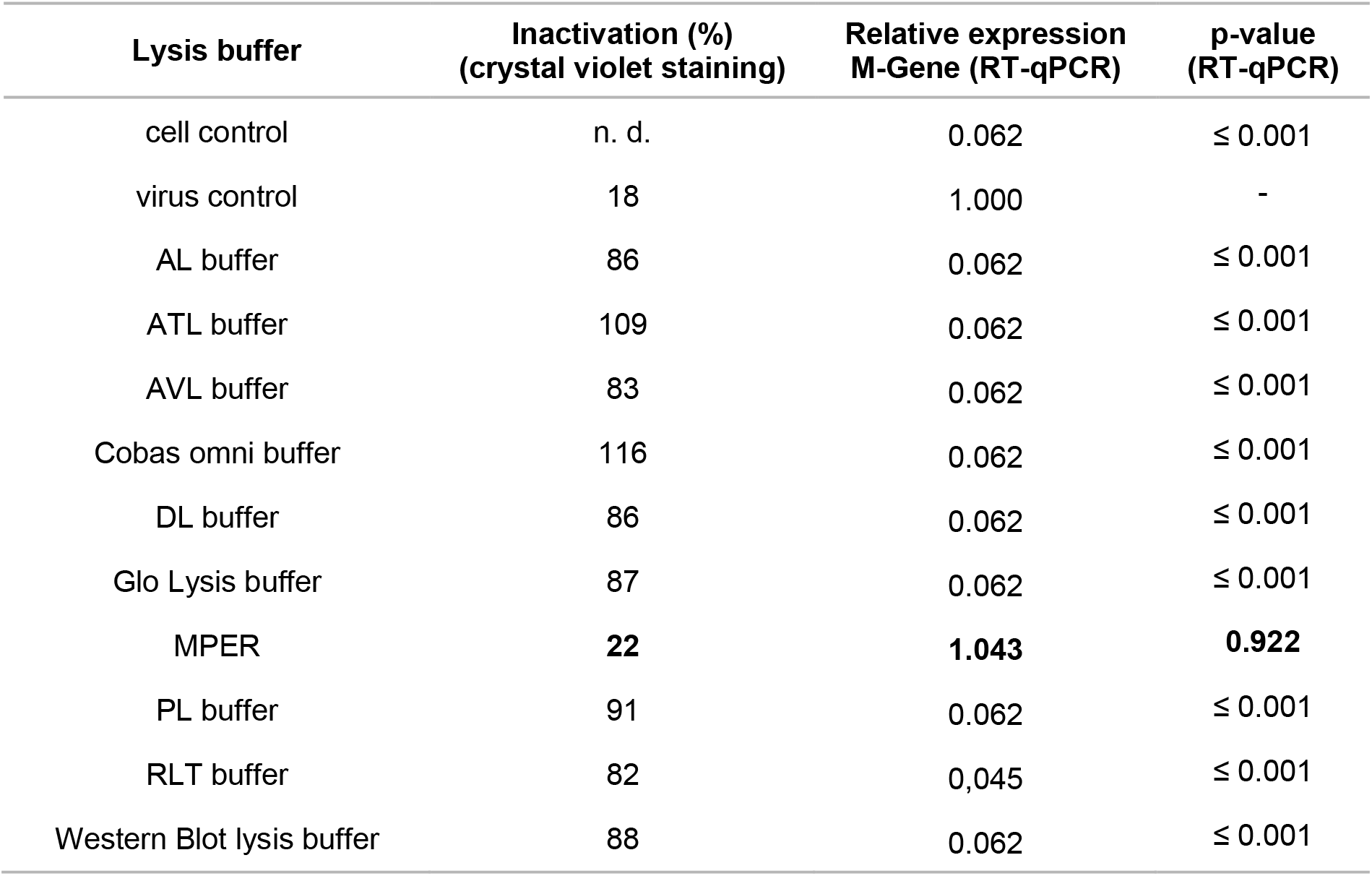
Inactivation of SARS-CoV-2 by lysis buffers commonly used in laboratory settings. Percent inactivation relative to OD_570_ in Crystal Violet staining, normalized to cell control (without virus). RT-qPCR data showing relative SARS-CoV-2 M gene expression for treated samples related to the virus control.

| Lysis buffer | Inactivation (%)<br>(crystal violet staining) | Relative expression<br>M-Gene (RT-qPCR) | p-value<br>(RT-qPCR) |
| --- | --- | --- | --- |
| cell control | n. d. | 0.062 | ≤ 0.001 |
| virus control | 18 | 1.000 | - |
| AL buffer | 86 | 0.062 | ≤ 0.001 |
| ATL buffer | 109 | 0.062 | ≤ 0.001 |
| AVL buffer | 83 | 0.062 | ≤ 0.001 |
| Cobas omni buffer | 116 | 0.062 | ≤ 0.001 |
| DL buffer | 86 | 0.062 | ≤ 0.001 |
| Glo Lysis buffer | 87 | 0.062 | ≤ 0.001 |
| MPER | **22** | **1.043** | **0.922** |
| PL buffer | 91 | 0.062 | ≤ 0.001 |
| RLT buffer | 82 | 0,045 | ≤ 0.001 |
| Western Blot lysis buffer | 88 | 0.062 | ≤ 0.001 |

**Figure 2.**
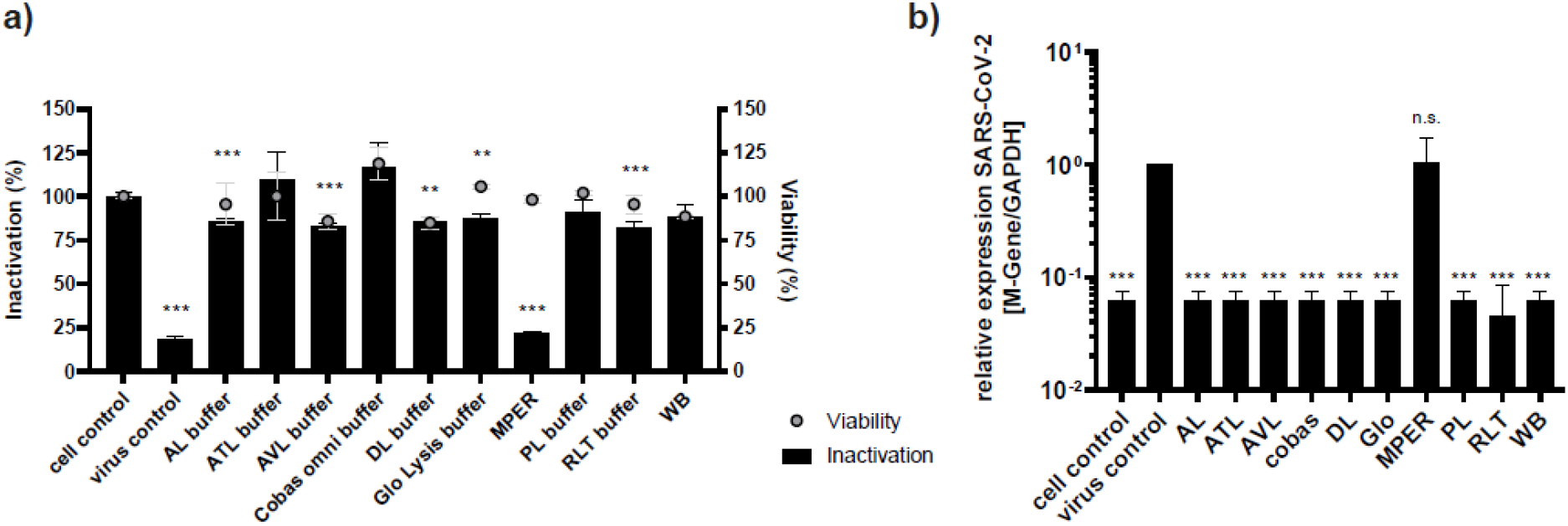
Inactivation of SARS-CoV-2 by lysis buffers commonly used in molecular biology laboratories. a) Crystal violet staining of Caco2 cells previously infected with virus pre-incubated with the indicated lysis buffer. Cell viability was determined by commercial RotiTest Vital Assay measuring OD_450_. Cells were incubated with a mixture of cell culture medium and the depicted lysis buffer in the given concentration used for virus inactivation. For the untreated control no viability assay was performed to avoid handling with infectious samples outside the safety cabinet. Samples were diluted 1:1000 to avoid cell toxicity. b) RT-qPCR analysis of intracellular RNA obtained from infected cells Caco2 cells showing the relative expression of SARS-CoV-2 RNA targeting M-Gene. Values were normalized to cellular GAPDH expression. Error bars indicate SD from the mean of representative experiment performed in triplicates. Virus control indicates SARS-CoV-2 infected cells without pre-incubation. Cell control indicates uninfected Caco2.

The IGEPAL CA-630 (MP Biomedicals Germany GmbH, Eschwege, Germany) containing direct lysis (DL) buffer was developed in order to avoid expensive and time-consuming RNA extraction procedure while concomitantly inactivating infectious samples prior to PCR analysis. DL buffer was found to efficiently inactivate SARS-CoV-2 (**Table 2**; **Supplementary Figure 1**). Next, we also tested lysis buffers that are commonly used to analyze protein samples and used a buffer for Western blot analysis, and Triton-X containing buffers, which are commonly used for luciferase-based assay. The latter have the critical property of maintaining protein function in order to allow enzymatic reactions with light emission. The buffers used therefore have a membrane-permeabilizing, but incompletely denaturing effect on cells. For Western blot lysis buffer and the commercially available Glo-Lysis buffer (Promega, Walldorf, Germany) as well as the PL buffer (lysis juice; p.j.k., Kleinblittersdorf, Germany) we were able to demonstrate complete inactivation (**Table 2**, **Supplementary Figure 1**). Importantly, Mammalian Protein Extraction Reagent (M-PER, ThermoFisher) was not able to inactivate SARS-CoV-2 (**Figure 2**, **Table 2**). As determined using crystal violet staining, only 22% of the cells were inactivated, which was comparable to the untreated control (18%). This data was confirmed by RT-qPCR since the relative SARS-CoV-2 M gene expression for infected and M-PER treated samples (1.043) was not statistically different from the virus control.

These findings highlight the fact that careful evaluation of the used inactivation methods are required and that additional inactivation steps might be necessary to prepare the samples for safe processing.

### 3.3 Physical Inactivation effectively eliminates infectious SARS-CoV-2

Since not all lysis buffers were able to completely inactivate SARS-CoV-2, we have further tested physical and chemical inactivation methods that can be applied subsequently. First, we evaluated the effects of heat inactivation on SARS-CoV-2 infectivity and heated cell culture supernatants at different temperatures with different incubation times. To monitor the temperature of the liquid that has to be inactivated, we generated heat curves (**Supplementary Figure 2, Supplementary Tables 1–2**). Subsequently, the treated supernatants were used to infect Caco2 cells. After 48 h cell culture supernatants were harvested and subjected to RT-qPCR to evaluate viral outgrowth (**Table 3**). We found that heat inactivation by placing the tube (500 μl in a 1.5 ml vessel) from ambient temperature (approx. 23°C) to a pre-warmed heating block for 5 min at 56°C was not effective (0.15 log), while increasing the incubation time to 30 min was a highly effective method of inactivation (4.35 log). Already 5 minutes at 60°C drastically reduced the infectivity of the virus suspension by 4.35 log. Short incubation time of 1 min at 90°C also achieved a very effective 4.28 log reduction in SARS-CoV-2 viral load.

**Table 3.** Heat inactivation, log reduction. Quantification of infectious SARS-CoV-2 using microscopical CPE based TCID_50_..

| Temperature (°C) | Incubation time (min) | Virus titre (TCID<sub>50</sub>/ml [log10] ± SD) | Minimal reduction factor (log10) | p-value |
| --- | --- | --- | --- | --- |
| Viral Control |  | 7.79 ± 7.22 | 0.00 | - |
| 56 | 5 | 7.63 ± 7.05 | 0.15 | 0.185 |
|  | 30 | 3.44 ± 2.71 | 4.35 | 0.003 |
| 60 | 5 | 3.54 ± 2.67 | 4.25 | 0.003 |
|  | 30 | 3.79 ± 3.70 | 3.99 | 0.003 |
| 90 | 1 | 3.95 ± 3.99 | 4.28 | 0.003 |
|  | 5 | 4.08 ± 3.99 | 3.71 | 0.003 |

Next, we evaluated chemical disinfectant and fixation solutions and treated virus supernatants in cell culture media in 1:10 dilution to the testing compound for 10 minutes at ambient temperature. Acetone/methanol (40:60), ethanol (70%), and paraformaldehyde (PFA, 3%) were found suitable to completely inactivate the virus (***Table 4***). However, ethanol/PBS (50%, 1:1) did not inactivate SARS-CoV-2, which was in line with previously published data (Kratzel et al., 2020b).

**Table 4.** Fixation of SARS-CoV-2. Virus load was determined via RT-qPCR targeting SARS-CoV-2 M gene with a detection limit of 3.3 copies/reaction (Log10). 4 log_10_ reduction was considered as antiviral activity.

| Fixation solution | Virus load* (copies/reaction [Log 10] ± SD) | Minimal reduction factor (Log10) | p-value |
| --- | --- | --- | --- |
| Viral Control | 7.63 ± 7.23 | 0.00 | - |
| Acetone/Methanol (40:60) | 3.34 ± 2.96 | 4.38 | 0.012 |
| 70% Ethanol | 3.65 ± 3.50 | 4.02 | 0.012 |
| 50% Ethanol/PBS (1:1) | 7.69 ± 6.85 | 0.00 | 0.568 |
| 3% Paraformaldehyde | 3.82 ± 3.05 | 3.81 | 0.012 |

Finally, we investigated the influence of UV light on the stability of SARS-CoV-2. For comparison, we included two different strains of SARS-CoV (from 2003) and also H1N1 Influenza A virus (IAV) as a representative of a seasonally recurring respiratory pathogen. To mimic daily laboratory decontamination routines, we exposed SARS-CoV-2 contaminated samples in a biological safety cabinet to UV light and irradiated for 15-60 min. We found that 15 min were already highly efficient and 30 min as well as 60 min completely inactivated SARS-CoV-2 infectivity. In addition, we used two different UV light sources with a discharge lamp (low pressure radiation chamber) or a highly potent LED UV light to inactivate contaminated surfaces. We compared different exposure times ranging from 2 – 210 seconds for UV-C discharge lamp and 0.5 −3.5 seconds for UV-C LED with fixed distances of 20 to 40 mm to the light source, respectively. Using UV light from the discharge lamp, 2 seconds exposure time resulted in a minimal reduction factor of 3.5 log while after 4 seconds no infectious SARS-CoV-2 was detected in the viral outgrowth assay (**Figure 3**, **Table 5**). One second UV-C LED was sufficient to reduce viral infectivity by 3.9 log. 3.5 seconds completely reduced viral titers >4 log.

**Table 5.** Inactivation of SARS-CoV-2 by UV-irradiation. Quantification of SARS-CoV-2 using microscopical CPE based TCID_50_. After exposure to UV-C light for the indicated exposure time SARS-CoV-2 was inoculated on Caco2 cells. Detection limit: 3.2 ×10^0^ TCID_50_/mL. 4 log_10_ reduction was considered as antiviral activity.

| Light source | Exposure time (s) | Virus titer (TCID<sub>50</sub>/ml [log10] ± SD) | Minimal reduction factor (Log10) | p-value |
| --- | --- | --- | --- | --- |
| UV-C discharge lamp | Viral control | 4.51 ± | - | - |
|  | 2 | 1.0 ± | 3.5 | ≤ 0.001 |
|  | 4 | 0.5 ± | 4.0 | ≤ 0.001 |
|  | 15 | 0.5 ± | 4.0 | 0.035 |
|  | 35 | 0.5 ± | 4.0 | 0.035 |
|  | 210 | 0.5 ± | 4.0 | 0.035 |
| UV-C LED | Viral control | 4.73 ± | - | - |
|  | 0.5 sec | 3.01 ± | 1.7 | 0.179 |
|  | 1.0 sec | 0.8 ± | 3.9 | 0.174 |
|  | 3.5 sec | 0.5 ± | 4.0 | 0.175 |

**Figure 3.**
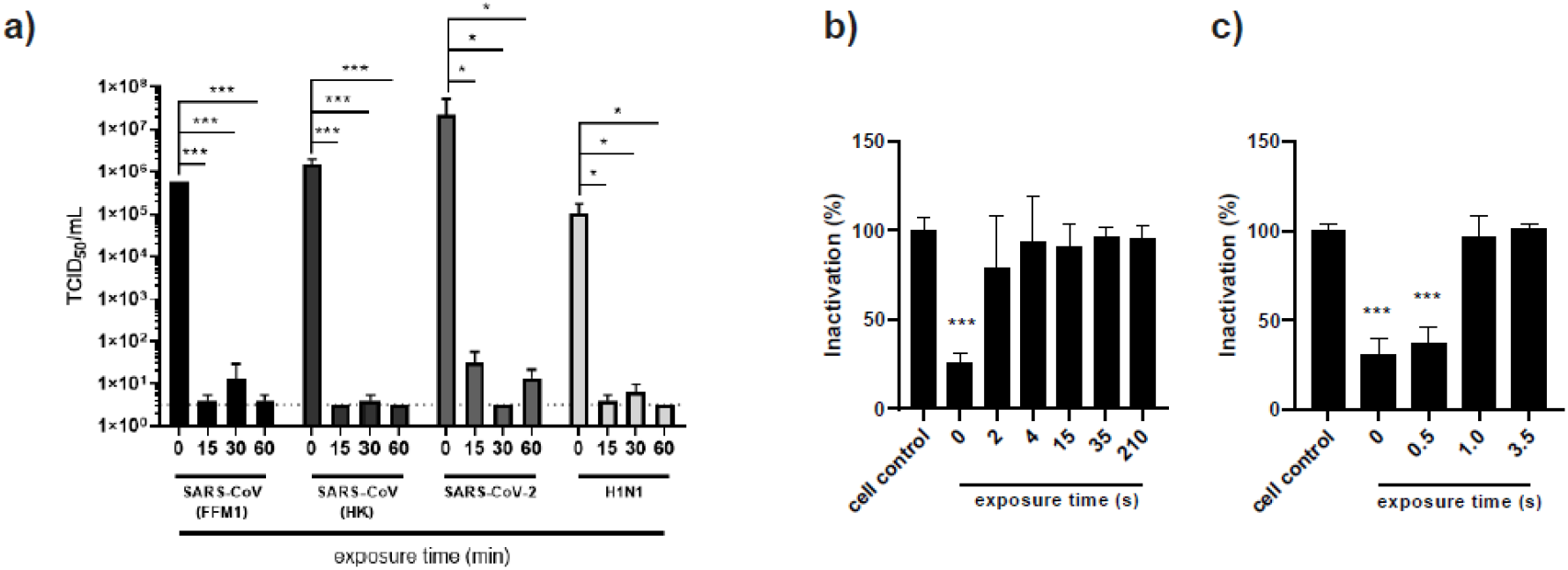
Inactivation of SARS-CoV-2 using UV-C light. **a)** Quantification of SARS-CoV-2 using microscopical CPE based TCID_50_ in Caco2 cells. After exposure to UV-C light in the safety cabinet for the indicated exposure time SARS-CoV-2 was used to infect CaCo2 cells. SARS-CoV-strains FFM1 (Drosten et al., 2003) and Hongkong (HK), SARS-CoV-2 strain FFM1 (Hoehl et al., 2020; Toptan et al., 2020), H1N1 (PR8). **b)** and **c)** Cristal violet staining of CaCo2 cells infected with UV-irradiated SARS-CoV-2. UV-C discharge lamp (b) and UV-C LED (c) were used as light source for the indicated exposure time. s, seconds. Error bars indicate SD from the mean of three independent experiments.

These data demonstrate that UV-light exposure is a suitable method for complete inactivation of SARS-CoV-2 contaminated material. In particular, irradiation with UV-C LED light might also represent a fast and reliable inactivation method for contaminated material and surfaces also in laboratory settings.

## 4. Discussion

Enveloped viruses are less resistant to chemical and physical environmental influences compared to non-enveloped viruses (Geller, Varbanov, and Duval, 2012; Kramer, Schwebke, and Kampf, 2006). Depending on air temperature and relative humidity (Casanova et al., 2010) coronaviruses are stable for up to 9 days on inanimate surfaces like metal, glass or plastic (Kampf et al., 2020b; van Doremalen, Bushmaker, and Munster, 2013). In this study mid-term and long term storages revealed a high stability of the virus during liquid storage over several weeks/months. We have found that even after months of storage conditions at 4°C, we found considerable quantities of infectious virus. This suggests that laboratories must be absolutely certain that their inactivation methods used for subsequent research purposes will also provide sufficient inactivation. SARS-CoV-2 can be efficiently inactivated by surface disinfection procedures (Kampf et al., 2020b), however, several research methods in molecular biology require functional proteins for enzymatically catalyzed reactions or for the maintenance of secondary and tertiary structures. In this manuscript, we tested several lysis buffers and other chemical and physical inactivation methods that would allow downstream analysis of SARS-CoV-2 infection outside of the BSL3 laboratory. Among several lysis buffers we found the proprietary detergent M-PER lysis buffer (Thermo Fisher) to fail in the ability to inactivate SARS-CoV-2.

In agreement with previously published studies we found that most inactivation methods were effective. In particular, our results were in line with studies that showed high temperature (Kratzel et al., 2020a), alcohols (Kratzel et al., 2020b), UV irradiation (Duan et al., 2003), and lysis by detergents are effective in inactivate corona virus (Rabenau et al., 2005a; Rabenau et al., 2005b). In particular, UV irradiation of infected cell culture samples was very effective in our study, but varied between the methods. Our results were partly consistent with previously published results recently reviewed by Derraik and colleagues (Derraik et al., 2020). In a study by Darnell et al., an experimental setup similar to our setup was performed, and revealed that already 1 min UV-C (distance from source 3 cm) irradiation led to a decrease in infectious SARS-CoV titers and that 6 min irradiation led to an almost complete inactivation (Darnell et al., 2004). After 15 minutes of UVC irradiation no detection of infectious viruses was possible.

These data are consistent with our observation that after 15 minutes UV-C irradiation in a sterile workbench more than 4-log loss in infectivity was observed for SARS-CoV-2. After irradiation with a highly potent UV-C source after 35 seconds and with a UV-C LED even after 1 second the vast majority of viruses were already inactivated.

The usual radiation takes place within the safety cabinets after work. Here, 15 minutes were already highly effective to eliminate infectious virus. In most laboratories, an exposure time of 30 minutes is suggested after infectious work. Thus, after correct use, inactivation of SARS-CoV-2 can be assumed, however, the physical lifespan of the light source must not be exceeded in order to generate sufficient emission. For fast inactivation of small surfaces, we evaluated two different UV-C light sources and found that UV-C discharge lamp with a radiation duration of 4 seconds was sufficient to inactivate SARS-CoV-2. The use of the high-potency UV-C LED light already results after 1 seconds. Although, all tested UV light source were UV-C rays the distance between light source and sample plays a critical role to define irradiation time. Furthermore, it has to take in account that lifespan of conventional UV lamps used in safety cabinets ranges between 6000 to 8000 hours. Therefore, intensity and effectiveness of the UV light is decreasing over time and when reaching maximum of expected useful life. In conclusion, irradiation with UV-C LEDs thus represents a highly effective inactivation method for contaminated surfaces.

In comparison to the Triton-X-containing lysis buffers used for enzyme based assays (e.g. luciferase based measurements), we found that the commercially available M-PER buffer (Thermo Scientific) was not sufficient to inactivate SARS-CoV-2. M-PER is a proprietary detergent in 25 mM bicine buffer (pH 7.6) used for enzyme based reporter protein assays and immunoassays as well as protein purification. While bicine is an organic compound used as a buffering agent the exact composition of M-PER is not publicly known. However, according to the manufacturer, M-PER reagent has been tested on different cell lines showing complete lysis of adherent cells. Even if the manufacturers describe an efficient and fast cell lysis, the virus was able to withstand the mild detergent in our experiments. In line with previously published data on SARS-CoV (Rabenau et al., 2005a), also fixation with PBS/ethanol (EtOH 50%; 1:1) over 5 min was sufficient to eliminate infectivity.

This study has limitations that need to be considered. To maintain real-lab-settings, we performed the stability evaluation with daily used virus stocks thawed for routine infectious experiments. Since the stocks in use had different lifetimes, longer standing times at ambient temperature could possibly have influenced viral stability. Hence we quantitatively cannot compare the samples with each other and also statistical evaluation was not applicable. Far more replicates under more controlled conditions would be necessary here. Nonetheless, even after 23 weeks we still found high-titer virus that has to be adequately inactivated by appropriate lysis buffer. These data are therefore important as a derivation for the inactivation tests. A further limitation of this study is based on the fact that inactivation efficiency largely depends on the initial virus load and the reaction volume. Especially in the case of heat inactivation, the heating time of the respective medium must be considered (**Supplementary Figure 1, Supplementary Tables 1–2**). In addition, the higher the virus titer, the longer the inactivation time is necessary for complete inactivation.

SARS-CoV-2 is able to replicate in susceptible cells lines like Caco2 and Vero cells both yielding high viral titers in the cell culture supernatants. The main difference is the ability to form a CPE, which is definitely more pronounced in Caco2 cells (Hoehl et al., 2020). However, in this study we did not match results obtained from different experiments with dissimilar cell lines but rather compare heat inactivation conditions in a defined setup using a specific cell line. Since the respective cell line was used for readout purpose, only the reduction factor which mirrors the heat inactivation efficiency is relevant.

In conclusion, inactivation methods including alcohols (except PBS/EtOH 1:1), acetone-methanol mixture, PFA, UV-light and heat inactivation efficiently inactivated SARS-CoV-2. Most but not all research laboratory used lysis buffers were suitable to inactivate SARS-CoV-2. However, since M-PER was not able to inactivate SARS-CoV-2, research buffers, especially if they are non-denaturating, must be carefully evaluated to protect laboratory personnel and must be substituted to maintain laboratory safety.

## 5 Acknowledgements

The authors thank Christiane Pallas and Leon Hildebrand for excellent technical assistance. Inactivation experiments were performed in cooperation with the Hönle AG who provided the UVA-Cube and the UV-C LED light source for testing. We are thankful for the numerous donations to the Goethe-Corona-Fond and for the support of our SARS-CoV-2 research. M.W. was supported by the Deutsche Forschungsgemeinschaft (DFG, WI 5086/1–1). All authors have read and agreed to the published version of the manuscript.

## Supplementary Figures

**Supplementary Figure 1.**
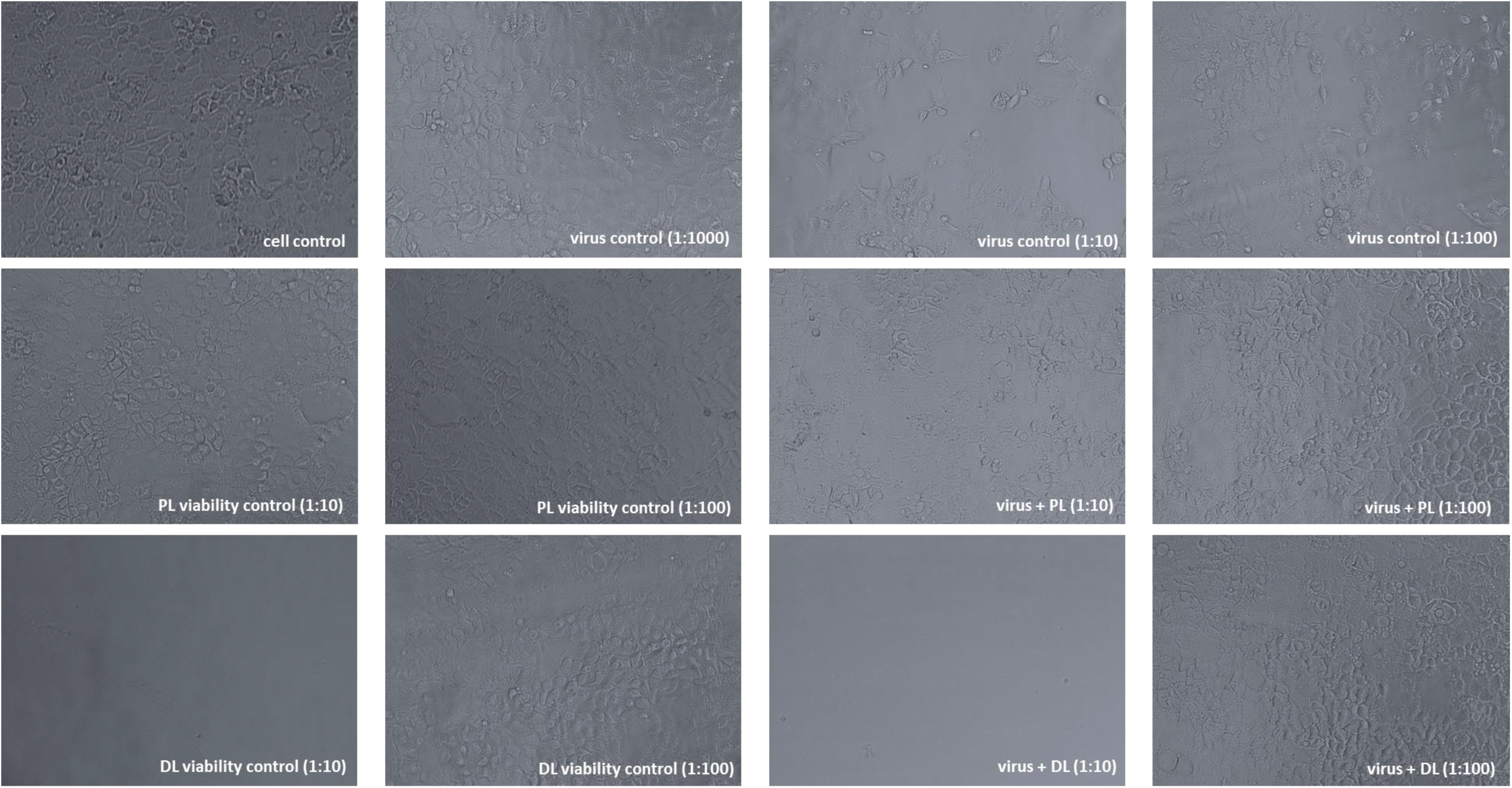
SARS-CoV-2 inactivation with Triton-containing lysis buffer (PL) and IGEPAL-630 containing buffer (DL). Crystal violet staining of CaCo2 cells infected with SARS-CoV-2 previously inactivated with the indicated buffer. Virus and lysis buffer (1:1) were incubated for 15 min at ambient temperature. Samples were diluted as indicated in brackets to avoid cell toxicity

**Supplementary Figure 2.**
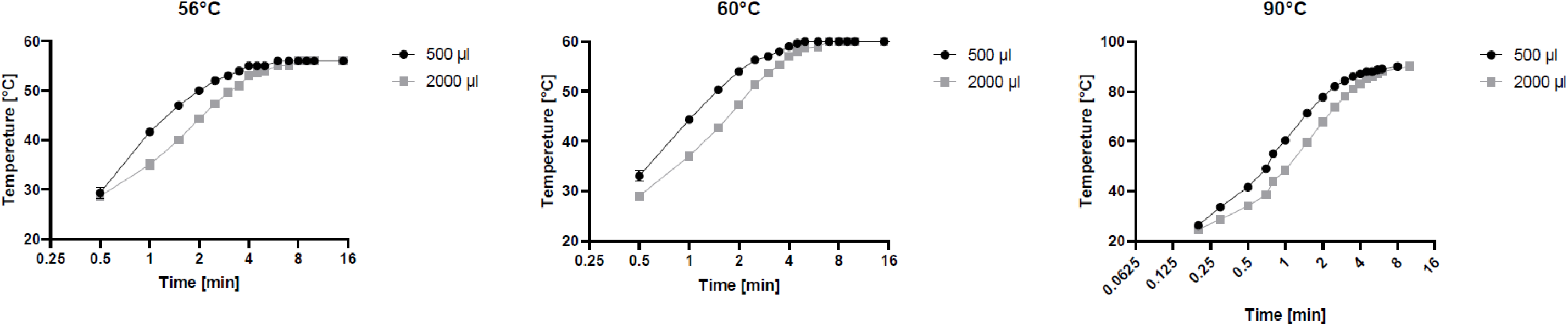
Heating curve of liquids in common laboratory reaction vessels. In order to minimize evaporation, 500 μl (black circles and line) or 2000 μl (grey squares and line) glycerol in a 1.5 ml or 2 ml reaction vessel, respectively, were placed in a preheated heating block. The increase in temperature was monitored by a thermometer immersed in glycerol. Mean values (n=3) were rounded up to the nearest integer. Error bars indicate standard deviation.

## Supplementary Tables

**Supplementary Table 1:** Heating curve of liquids in common laboratory 1.5 and 2 ml reaction vessels placed in a heat block pre-warmed to 56°C and 60°C, respectively. The increase in temperature was monitored by a thermometer immersed in 500 μl (1.5 ml vessel) and 2000 μl (2 ml vessel) glycerol, respectively. Experiments were performed in triplicates. Mean values were rounded up to the nearest integer.

| Time (min) | Temperature (°C)<br>500 µl at 56°C | Temperature (°C)<br>2000 µl at 56°C | Temperature (°C)<br>500 µl at 60°C | Temperature (°C)<br>2000 µl at 60°C |
| --- | --- | --- | --- | --- |
| 0 | 23 | 23 | 23 | 23 |
| 0.5 | 29 | 29 | 33 | 29 |
| 1 | 42 | 35 | 44 | 37 |
| 1.5 | 47 | 40 | 50 | 43 |
| 2 | 50 | 44 | 54 | 47 |
| 2.5 | 52 | 47 | 56 | 51 |
| 3 | 53 | 50 | 57 | 54 |
| 3.5 | 54 | 51 | 58 | 55 |
| 4 | 55 | 53 | 59 | 57 |
| 4.5 | 55 | 54 | 60 | 58 |
| 5 | 55 | 54 | 60 | 59 |
| 6 | 56 | 55 | 60 | 59 |
| 7 | 56 | 55 | 60 | 60 |
| 8 | 56 | 56 | 60 | 60 |
| 9 | 56 | 56 | 60 | 60 |
| 10 | 56 | 56 | 60 | 60 |
| 15 | 56 | 56 | 60 | 60 |
| 20 | 56 | 56 | 60 | 60 |
| 25 | 56 | 56 | 60 | 60 |
| 30 | 56 | 56 | 60 | 60 |
| 35 | 56 | 56 | 60 | 60 |

**Supplementary Table 2:** Heating curve of liquids in common laboratory 1.5 and 2 ml reaction vessels placed in a heat block pre-warmed to 90°C. The increase in temperature was monitored by a thermometer immersed in 500 μl (1.5 ml vessel) and 2000 μl (2 ml vessel) glycerol, respectively. Experiment was performed in triplicates. Mean values were rounded up to the nearest integer.

| Time (min) | Temperature (°C)<br>500 µl at 90°C | Temperature (°C)<br>2000 µl at 90°C |
| --- | --- | --- |
| 0.0 | 23 | 23 |
| 0.2 | 26 | 25 |
| 0.3 | 34 | 29 |
| 0.5 | 42 | 34 |
| 0.7 | 49 | 39 |
| 0.8 | 55 | 44 |
| 1.0 | 60 | 48 |
| 1.5 | 71 | 60 |
| 2.0 | 78 | 68 |
| 2.5 | 82 | 74 |
| 3.0 | 84 | 78 |
| 3.5 | 86 | 81 |
| 4.0 | 87 | 83 |
| 4.5 | 88 | 85 |
| 5.0 | 88 | 86 |
| 5.5 | 89 | 87 |
| 6.0 | 89 | 88 |
| 8.0 | 90 | - |
| 10.0 | - | 90 |

